# CircMarker: A Fast and Accurate Algorithm for Circular RNA Detection

**DOI:** 10.1101/134411

**Authors:** Xin Li, Chong Chu, Jingwen Pei, Ion Măndoiu, Yufeng Wu

## Abstract

While RNA is often created from linear splicing during transcription, recent studies have found that non-canonical splicing sometimes occurs. Non-canonical splicing joins 3’ and 5’ and forms the so-called circular RNA. It is now believed that circular RNA plays important biological roles such as affecting susceptibility in some diseases. within these few years, several experimental methods have been developed to enrich circular RNA while degrade linear RNA. Although several useful software tools for circRNA detection have been developed as well, these tools may miss many circular RNA. Also, existing tools are slow for large data because those tools often depend on reads mapping. In this paper, we present a new computational approach, named CircMarker, based on k-mers rather than reads mapping for circular RNA detection. CircMarker takes advantage of transcriptome annotation files to create k-mer table for circular RNA detection. Empirical results show that CircMarker outperforms existing tools in circular RNA detection on accuracy and efficiency in many simulated and real datasets. CircMarker can be downloaded from https://github.com/lxwgcool/CircMarker.

## 1 Introduction

In most eukaryotic genes, coding regions (exons) are separated from noncoding regions (introns). During the process of RNA splicing, introns are removed and exons are joined to form a contiguous coding sequence called messenger ribonucleic acid (mRNA). This “mature” mRNA is ready for translation, and those contiguous coding sequences are called transcription [2]. Splicing often occurs in a linear way, which generates the so-called linear RNA. Recent studies show that sometimes circular RNA may be generated during transcription [14]. Circular RNA (or circRNA) is a type of RNA which forms a covalently closed continuous loop. That is, the 3’ and 5’ ends normally present in an RNA molecule are joined together [1]. This feature leads to numerous properties of circular RNAs [15]. However, since the amount of circular RNA is often much lower than linear RNA, circular RNA has not been thoroughly studied until recently. During the past several years, several papers report that circular RNA may be associated with diseases and traits. The increasing number of circular RNAs have been identified recently [7, 6].

Since circular RNAs do not have 5’ or 3’ ends, they are resistant to exonuclease-mediated degradation and are presumably more stable than most linear RNAs in cells. Based on this feature, some benchmark experimental methods have been developed to degrade the linear RNA while enrich the circular RNA. For example, one method is treating samples with RNase R, an enzyme which degrades linear RNAs but not circular RNAs. This treatment can enrich circular RNAs [9]. Often circular RNA comes with the splicing signals of “AG” or “AC” as starting while “GT” or “CT” as ending for direction “+” and “-” respectively [8].

Computational tools for circular RNA detection have been developed. Currently, there are several existing tools for circular RNA detection, such as Find circ [13], CIRCexplorer [16] and CIRI [3]. Find circ is one of the first tools for circular RNA detection. Since it is difficult to map the joint position of circular splicing back to the reference genome, Find circ tries to collect all un-mapped reads based on reads mapping results from Bowtie [11]. Then, all unmapped reads are converted to new short reads by combining the head parts and the tail parts of current reads together. Then Find circ maps the new short reads back to the reference. CIRCexplorer performs reads mapping using Bowtie and TopHat. The main idea is using the concept of fusion gene to detect circular RNA. First, CIRCexplorer tries to find out the un-mapped reads. Then, those un-mapped reads are mapped back to the reference using TopHat-Fusion [10] to detect potential circular RNA candidates with the back-spliced junction reads. CIRI uses BWA for reads mapping, trying to find circular RNA by analyzing CIGAR signatures in the SAM file. Some of these tools such as CIRCexplorer depend on transcriptome annotation, while others support de novo circular RNA detection, such as Find circ.

All of these methods mentioned above depend on reads mapping. These mapping based methods have some inherent issues. The first issue is computational efficiency: the existing tools use BWA, Bowtie or TopHat for reads mapping. Although BWA and Bowtie are widely used in sequence analysis, reads mapping is still time-consuming for circular RNA detection. This is because reads mapping tries to map every read, even when the read is not relevant for circular RNA detection. In addition, since some new short sequences may be created in the middle by circRNA detection tools for the second round mapping, reads mapping can become very slow when TopHat-fusion is used, due to the length of sequences. Moreover, these tools may miss circular RNA in some cases due to errors in reads mapping. For example, some reads related to circular RNA may be un-mapped due to reads error.

In this paper, we develop a new computational method, called CircMarker, for circular RNA detection. The objective of CircMarker is finding the presence of circular RNA (in particular the join of two known exons). CircMarker doesn’t reconstruct the complete sequence of circular RNA. The key idea of CircMarker is that it doesn’t rely on reads mapping. Instead, CircMarker analyzes short sequence segments, called k-mers, for circular RNA detection. The main advantage of using k-mers is efficiency: finding k-mers from reads is much faster than reads mapping. Another advantage is that k-mer tolerates more errors in reads and carries useful information about the presence of circular RNA, which may be missed by reads mapping. Empirical results show that CircMarker is more accurate than (or as accurate as) existing methods on simulated and real datasets in calling circular RNA. CircMaker runs much faster than existing methods.

## 2 Method

**High-level approach**. The overall approach of CircMarker is shown in Figure 1. CircMarker is based on analyzing k-mers in the sequence reads. That is, CircMarker doesn’t perform reads mapping. CircMarker only considers the circular RNA which comes from the exons identified by annotation file. We do not consider de novo circular RNA cases. CircMarker uses three types of inputs, including the reference genome, the transcription annotation file and sequence reads. Note that all circular splicing that we consider here occurs at the boundary of exons identified by the given annotation file. CircMarker first processes the annotation file and the reference genome. It extracts and stores all k-mers that are located near the exon boundaries. To speed up, CircMarker first performs a fast check to find the reads that are likely to be relevant for circular RNA detection. Then it processes each read and compares k-mers in the read with the stored k-mers to identify circular RNA based on the signatures from circular RNA. When two k-mers from a single read are *out of order* with regard to the reference, CircMarker considers this as an evidence for the existence of circular RNA. This is illustrated in Figure 1.

**Fig. 1.**
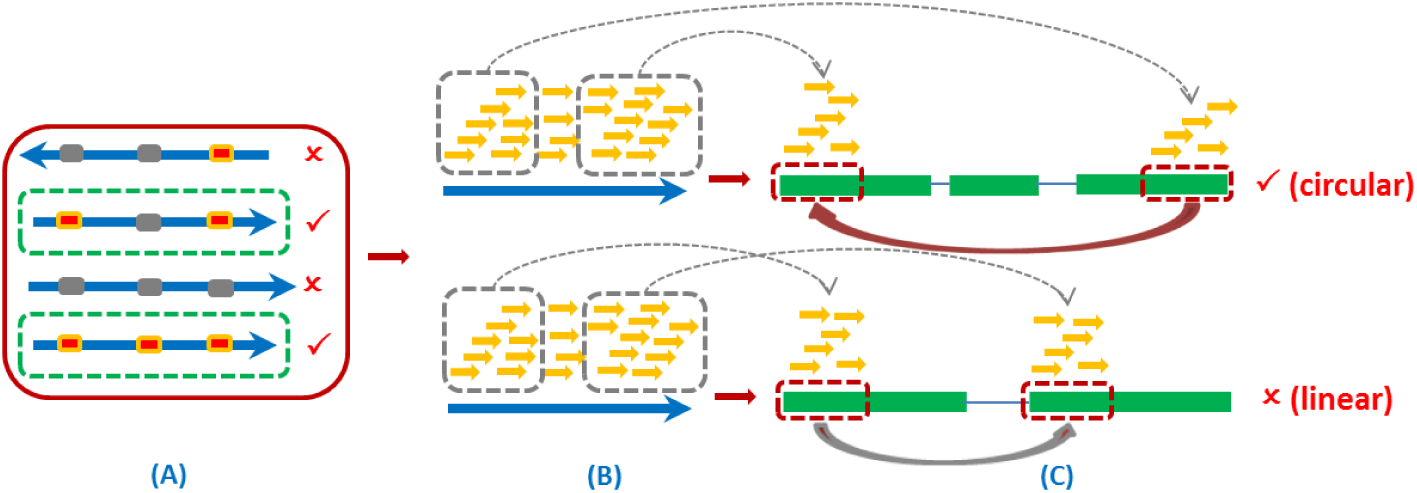
The procedure of the circular RNA detection. (A) A fast check for finding circular RNA relevant reads by sampling. The blue arrow stands for reads and the dots within it present the sparsely sampled k-mers. Gray dot: k-mer with no hit in the k-mer table from annotation. Red dot: k-mer which finds a hit in k-mer table. Reads inside green box: pass quick check. (B) Scanning k-mer sequentially from the beginning to the end for each read. Yellow arrow: k-mer. (C) Calling circular RNA using various criteria and filters. Green bar: exon along the reference. Two transcriptions are listed here. The upper: with 3 exons, and the red arrow identified a potential circular RNA. The lower: with 2 exons, and the gray arrow stands for linear RNA.

### 2.1 Processing the reference genome and annotations

CircMarker creates a table for storing the k-mers within the reference genome that are near the exon boundaries as specified by the annotations. The k-mer table is designed to be space-efficient. We only record the following five types of information for each k-mer, including chromosome index, gene index, transcript index, exon index and the part tag as shown in Figure 2. The “part tag” specifies whether the k-mer comes from the head (i.e. beginning) part or the tail (i.e. ending) part of the exon. Due to the relative small ranges of index, a record on a k-mer only needs eight bytes. We call it the de novo position. One k-mer may contain a group of de novo positions. 32 bits integer is used to store the information of a k-mer, which means the maximum length of a k-mer should be shorter than 16 bp, and all k-mers which contain invalid letters such as “N” are discarded.

**Fig. 2.**
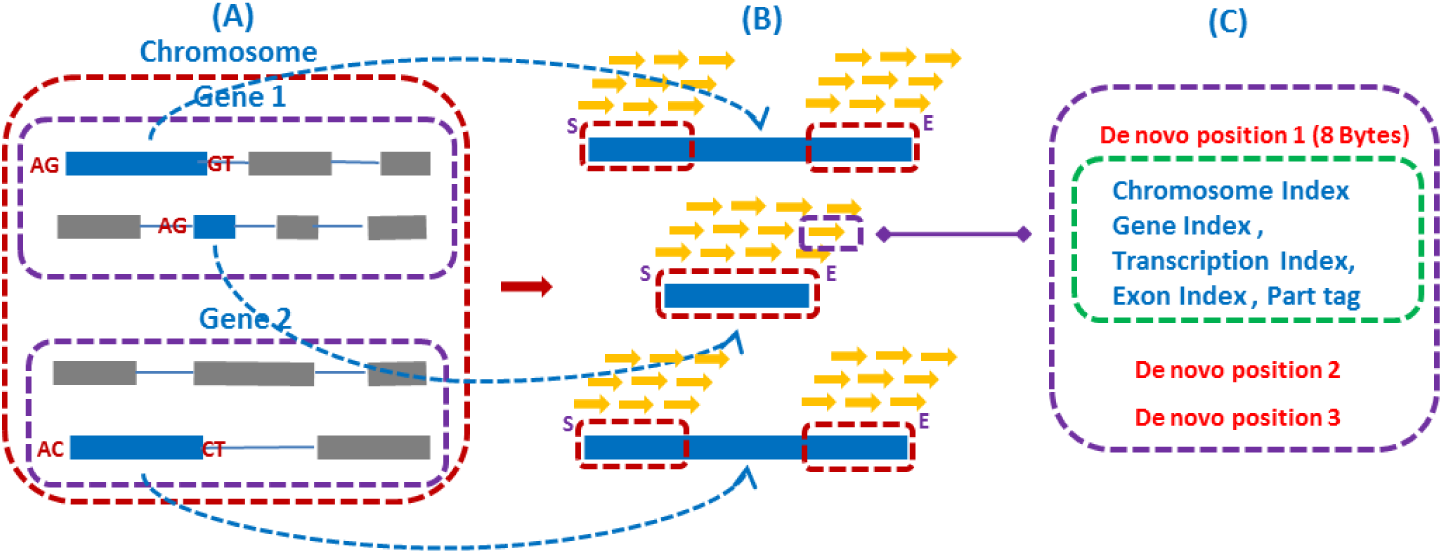
Collecting k-mers from annotated exons in reference genome. These k-mers are stored in a table that will be used to compare with the reads. (A) Annotations from one chromosome. Blue bars stand for valid exons while gray bars stand for the invalid ones which do not contain any circular splicing signal. (B) Extracting K-mer. Yellow arrows stand for the k-mers from the boundary (red frame) of each exon. All of k-mers from short exon will be considered (the second case). “S” or “E” is the value of part tag. (C) Showing all de novo positions of the k-mer with purple box in B). Green box: all types of information in one de novo position.

When extracting k-mers from annotated exons in the reference genome, we only consider the exons with circular splicing signal in either head or tail part. And we only consider the k-mers which come from the left and right boundaries of the exon. The length of the boundary region *L*_*B*_ is defined as below:

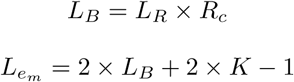

Here *K* is the length of the k-mer. *L*_*R*_ is the length of reads, and *R*_*c*_ represents the percentage of reads that should be covered in each boundary. Since we expect more than half reads to be considered, we set the default value of *R*_*c*_ as 30% (2 * 30% = 60% *>* 50%). If 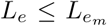, we use the whole exon to create k-mer and set the part tag as “S” if it located in the first half part and “E” for the second half. Otherwise, we use the head boundary and tail boundary of current exon to extract k-mers and set the part tag to “S” or “E” respectively.

### 2.2 Processing sequence reads

Once k-mer table of the annotated exons is created, we now process each sequence read. Here we examine k-mers contained in a read and search for a match in the k-mer table. This way, we obtain the “hitting status”, which means which transcription can be hit by current reads. “hit exon” means the exon that is hit by the k-mer in the reads in k-mer table. Each reads may related to more than one hitting status, each hitting status contains at least one hit exon, and each hit exon should be supported by at least one k-mer. We scan all reads to check their hitting statuses. In order to skip irrelevant reads, we perform basic check initially by sampling eight k-mers from 10% to 80% position of the current read.

The read passes the sampling check only if at least two k-mers can find a hit in the k-mer table. If so, we examine all of k-mers from start to end, collecting all hitting statuses in this order. Since each hitting status is contributed by multiple k-mers, the “best hitting case” is defined as:

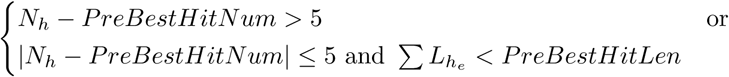

*N*_*h*_ specifies the number of k-mers which supports all of hit exons in one hitting status. The *PreBestHitNum* means the *N*_*h*_ of previous best hitting status, 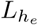 means the length of one hit exon in current hitting status, and *PreBestHitLen* means the summary length of the hit exons in previous best hitting status. If *N*_*h*_ is larger than *PreBestHitNum* + 5, the current hitting status will be set as the previous best hitting case, which means we prefer the hitting status with conditional larger number of k-mers supporters. Otherwise, the hitting status with the shorter summary length of the hit exon will be chosen. We set the previous best hitting status as the final best hitting case when all of hitting status been processed. Finally, the *N*_*h*_ of best hitting case should be at least 5. Otherwise it is discarded.

### 2.3 Filtering

The previous step identifies best hitting cases. Due to the inherent noise in the data (e.g. read errors and duplications), we perform the following filtering step to improve the accuracy. There are two main filtering procedures.

*Filtering procedure one.* The first filter procedure is checking the hitting number. The minimum hitting number 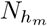 is defined as below:

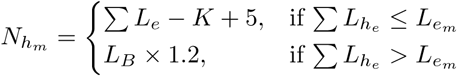

The key is that short exons should be fully covered by the reads more than one time. Otherwise, we need to ensure the reads to be within both boundaries of the hit exons. Any best hitting case is discarded if the *N*_*h*_ is smaller than 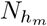.

*Filtering procedure two.* Based on the number of hit exons in best hitting case, we divide all cases into two types: the case of self-circular if the number of hit exons is equal to 1, and the regular-circular case otherwise.

For self-circular case, only the exon containing circular splicing signal in both sides will be considered. Then, the best hitting case will be considered as the self-circular RNA candidate if 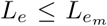. Otherwise, we collect the part tags from begin to end, and condense the tags which belong to the same exon based on the number of hitting. For example, we define *S*(*n*) as n continuous tag “S” in one exon (similar for *E*(*n*)). If we have *S*(1) and *E*(10) in one exon, then we condense them to *E*(10). This may help us to filter some random hits. We consider a candidate a valid self-circular RNA only if the number of tags after condensation is equal to 2 and the tags are arranged from E to S sequentially (i.e. going backward at the circular RNA join junction).

For the regular-circular case, the best hitting case will be considered only if it contains two exons. First, an exon will be skipped if its hitting time is at most 3 in order to remove some random hits. Then, we try to condense the tags. Here the method described in the self-circular case will be applied at first. After that, for the first exon we condense SE to E while condensing SE to S for the second exon. This condensation logic may solve the problem that some of exons are fully covered by current reads. The best hitting case will be kept only if the number of condensed tag in both exons is equal with 1 and the tags arranged from E to S sequentially.

### 2.4 Calling circular RNA

There are two cases for calling circular RNA: the self-circular case and the regular-circular case.

*Self-Circular RNA.* First, a self-circular RNA candidate will be discarded if the length of current exon is shorter than the read length while the *N*_*h*_ is smaller than *L*_*e*_− *K* + 1. Otherwise, the best hitting case will be considered to be a valid self-circular RNA candidate if it contains circular splicing signals in both sides.

*Regular-Circular RNA.* For the direction “+”, the candidate will be dropped if the exon index increases monotonically. Otherwise, we try to identify the breakpoint at the position of the first deceasing and set it to be the joint junction of circular RNA. We call the exon with large index as the head exon while another one as the tail exon. Based on this definition, the head exon is located in the later part of the reference, while tail exon is located in the earlier part, and the circle should connect the head exon back to the tail exon. The candidate will be viewed as a valid regular-circular RNA candidate only if the head exon and tail exon have the tail and head circular splicing signal respectively. We set the end position of head exon and the start position of tail exon as the position of this called regular-circular RNA.

For the direction “-”, the procedures is mostly the same as the direction “+”. The only difference is how to choose the joint junction. In this case, the candidate will be dropped if the exon index decreases monotonically. Otherwise we try to identify the breakpoint at the the first increasing and set it to be the joint junction of circular RNA. The exon with small index is viewed as the head exon while the big index exon is set as tail exon.

*Refining circular RNA candidates.* We count how many reads support each circular RNA candidate. Only the candidate with support number smaller than the predefined threshold will be viewed as the correct one. Since the maximum coverage of circular RNA is unknown in most cases, we set the default value to be 999 to allow all of valid circular RNA candidates.

## 3 Results

Since the study of circular RNA is still at an early stage, there is no widely accepted benchmark data for evaluating the circular RNA calling at present. Recently, there are some public circular RNA databases which collect different types of circular RNA from published papers. Some databases come with the recommended circular RNA detection tool, such as CircBase [5]. Others focus on collecting the relationship between circular RNA and diseases or traits, such as Circ2Traits [4].

In this paper, we use both simulated and real data to compare CircMarker with three existing tools, including CIRI, Find circ, and CIRCexplorer in terms of the number of called circular RNA, accuracy, consensus-based sensitivity, bias and running time. When comparing the genomic positions of circular joint junction, we allow up to five bp tolerance. Since CircMarker is based on k-mers and each chromosome has its own k-mer table, the running time can be reduced significantly by parallelization (i.e. running analysis on each chromosome in parallel).We compare the performance of these tools on the first three chromosomes individually. Because some existing tools do not support parallelization, we use a single core to run each program for circular RNA detection, and use 10 to 12 cores to run the reads mapping programs such as BWA, Bowtie and TopHat.

### 3.1 Simulated data

We first use simulated data for evaluation. To generate simulated data, we use the simulation script (called “CIRIsimulator.pl”) released by CIRI. The reference genome is the chromosome 1 in human genome (GRCh37). The annotation file is the version 18 (Ensembl 73). Two different cases are simulated as follows: (1) pair-end reads with 13,856,032 sequences, which roughly lead to 10X coverage for circular RNA and 100X coverage for linear RNA, and (2) pair-end reads contains with 9,400,036 sequences, which lead to us 50X coverage for both circular and linear RNA. The goal of the case 1 simulation is simulating the regular RNA-seq, while the case 2 focuses on the situation when the coverage of circular RNA is higher. The reads length is 101 bp and the insert size is 252 bp in both cases. The total number of simulated circular RNA in benchmark is 8,033 and 8,071 for those two cases respectively. Note that the true circular RNA is known in simulated data, which can be used in comparison. Since the coverage of circular RNA is known in simulated data, we set the “maximum support reads” equal with 10 and 50 in CircMarker respectively. We use the following three statistics for comparison: (1) hit number *N*_*h*_: the number of called circular RNA that are true, (2) accuracy: 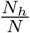 where *N* is the total number of called circular RNAs by a method, (3) running time.

The results of the four tools being compared are shown in Figure 3. Our results show that CircMarker outperforms the existing tools in terms of hit number, accuracy and running time. This is especially evident in case 1 (the left part in Figure 3), where CircMarker has fewer false positives and also calls more correct circular RNA than other tools. For case 2, the accuracy of CircMarker decreased to 32.04% from 70.90% in case 1. This is likely due to the week performance of the option “coverage filter”, for the similar coverage in both linear and circular RNA. Still, CircMarker is slightly more accurate than existing tools in this case. Moreover, CircMarker runs much faster than existing tools.

**Fig. 3.**
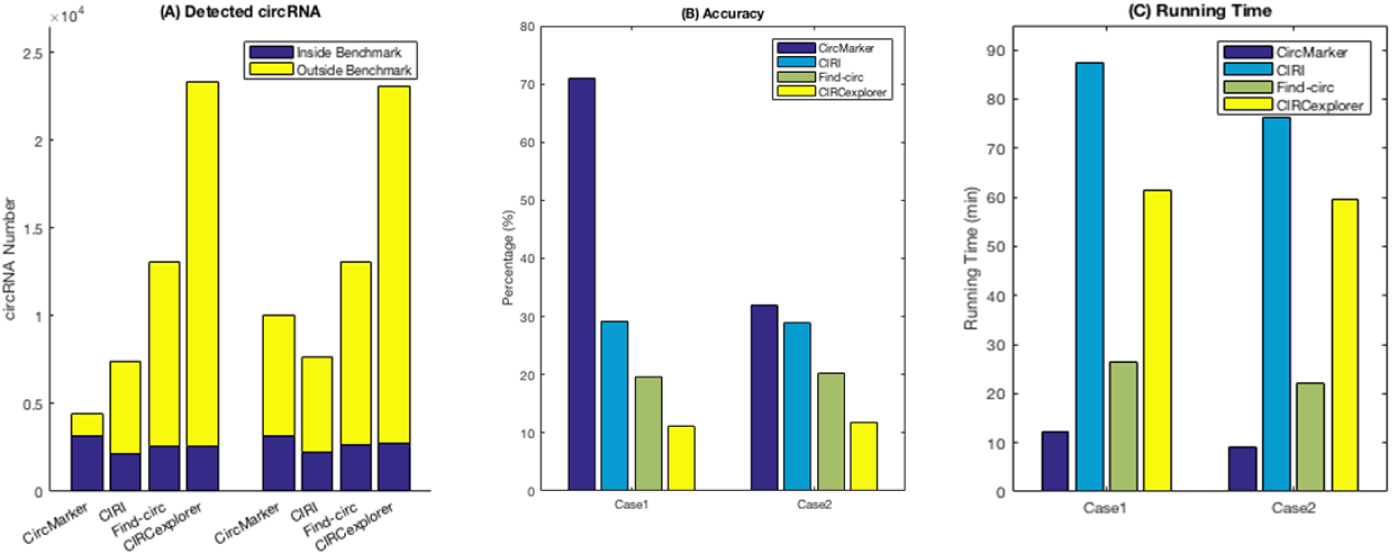
(A) The number of circular RNA called by each tool in case 1 (10X and 100X, the left cluster) and case 2 (50X&50X, the right cluster). Yellow bars: the number of un-hit (i.e. incorrectly called) circular RNA. Blue bars: the number of hit (i.e. correctly called) circular RNA. (B) The accuracy of each tool in cases 1 and 2. (C) The running time (in minutes) of each tool in both cases.

### 3.2 Real data

We use two types of real data to evaluate the performance of the four tools.

**Real RNase R treated sequence reads with public database information.** As described before, some public databases contain circular RNA called by published papers. In those papers, the authors usually only validate parts of the computationally detected circular RNA using biological experiments. The final result will be released only when the accuracy of those randomly chosen candidates meets certain standard. Therefore, we consider those released circular RNAs in these databases are reliable in this paper.

*Data Collection* We choose CircBase as the standard circular RNA database of *homo sapiens*. We use the circular RNAs recorded in this database as “benchmark”. The reference genome and annotation file come from homo sapiens GRCm37 version 75. The RNA-Seq reads are from SRR901967. These RNA-Seq reads are used to examine circular RNAs from RNase R treated poly(A)-/ribo-RNAs in human embryonic stem cells. There are total 41,342,095 single-end reads in this data.

We use the first three human chromosomes for comparison and use four statistics for comparison. (1) Hit number 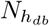: the number of circular RNA which has a matched circular RNA in the database. These matched circular RNA are called reliable circular RNA. (2) Intersection: the intersection of reliable circular RNA between CircMarker and other tools. This value could be used to evaluate the bias. (3) Reliability ratio: 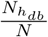. This measures the fraction of the number of matched circular RNA with regard to the total called ones *N*. (4) Running time. The best tool is expected to have large intersection with other tools (low bias), large number of reliable circular RNA with high reliability ratio.

The number of circular RNAs in CircBase from chromosome 1 to chromosome 3 is 9,142, 7,530 and 5,320 respectively. The results show that CircMarker finds more “benchmarked” circular RNAs and runs much faster than others (Figure 4 A, C). For the reliability ratio, there is a trade off with hit number. CIRI obtains the highest reliable ratio, but has the smallest hit number. The reliability ratio of CircMarker is similar to those of Find circ and CIRCexplorer (Figure 4 A). CircMarker has the largest hit number. In addition, CircMarker has the large intersection with the results from other tools in all three chromosomes, which means it has low bias (Figure 4 B). As a result, CircMarker outperform the other tools in this data.

**Fig. 4.**
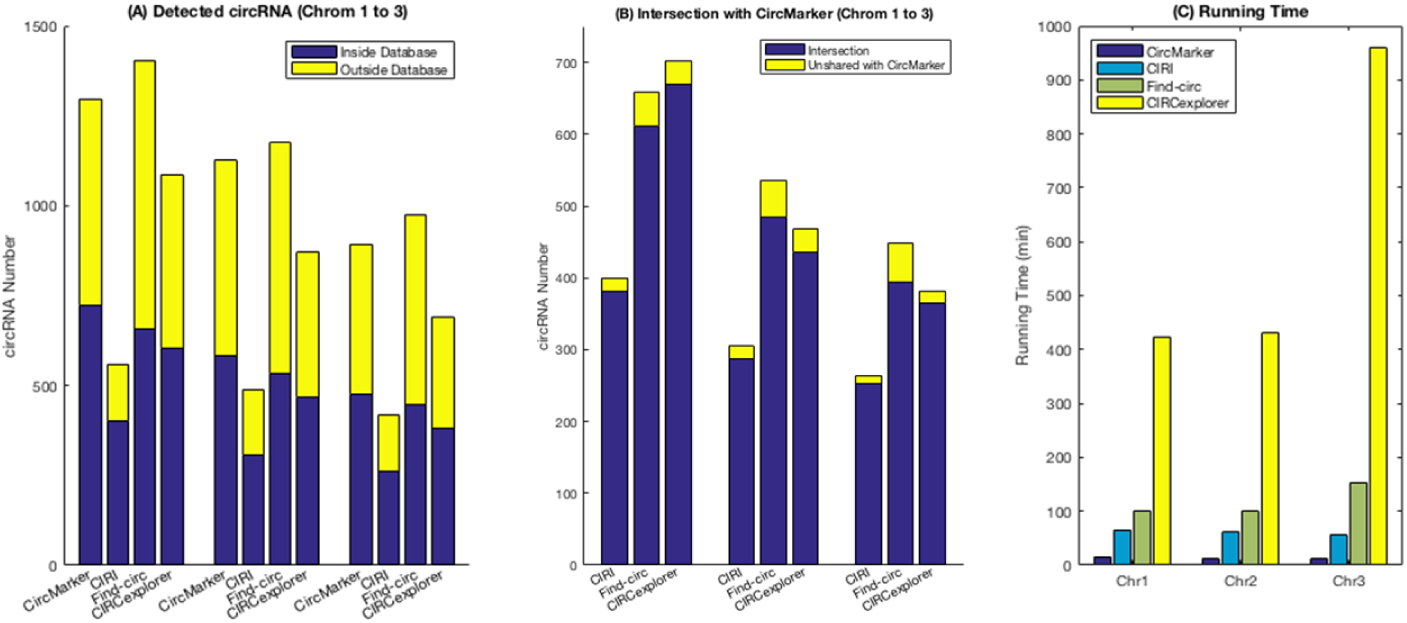
(A) The number of circular RNA called by each tool from chromosome 1 to chromosome 3. Yellow bars: the number of called circular RNAs which do not match with database. Blue bars: the number of circular RNAs that have matches in database. (B) Intersection: the number of circular RNAs in the intersection of reliable circular RNA 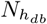 between CircMarker and other tools. High intersection ratio implies low bias. (C) The running time (in minutes) of each tool in all three chromosomes.

**Real RNase R Treated/Untreated Data** RNase R is an experimental technology that can break down linear RNA and enrich circular RNA. As a result, one popular way for validating a circular RNA detection tool is running the tool in two different types of reads: one from only rRNA eliminated sample (called untreated), and the other from RNase R treated sample. The circular RNA which can be found in both types of reads is considered to be reliable.

*Data Collection.* The reference genome and the annotation file are from *Mus Musculus GRCm38 Release79*. The RNase R treated reads are from SRR2219951 and the untreated reads are from SRR2185851. The library was prepared using the script Seq v2 Kit from Epicentre [12], and this data has been used to delineate the circular RNA complement of mouse brain at age 8 to 9 weeks. Both datasets contain pair-end reads, and SRR2219951 (treated) contains 44,661,952 sequences while SRR2185851 (untreated) contains 65,879,618 sequences.

We use the first three chromosomes of *Mus Musculus* with the two types of reads for comparison. We use the following three statistics. (1) Reliable circular RNA: the reliable circular RNA are from the intersection of called circular RNAs between the treated and untreated reads. Each tool reports its own reliable circular RNA from chromosomes 1 to 3. (2) Consensus-based sensitivity: we say a called circular RNA to be trusted if this circRNA is called by at least two tools. These trusted RNAs are considered to be “benchmark”. We collect these trusted circular RNA for each chromosome. Then, we calculate the intersection between the reliable circular RNA and the benchmark for each tool respectively from chromosome 1 to 3. The consensus-based sensitivity is calculated by: 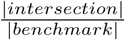. (3) Running time. Ideally, a circular RNA detection tool should obtain large number of reliable circRNA with high consensus-based sensitivity and fast running time in each chromosome.

The results are shown in Figure 5. CircMarker finds larger number of reliable circular RNA than others in all three chromosomes (Figure 5 A). The number of circRNAs in benchmark (i.e. trusted circRNA supported by at least two tools) is 322, 353 and 186 for chromosomes 1 to 3 respectively. One can see that CircMarker gets the largest number of reliable circular RNA in all three chromosomes. In addition, it gains the highest consensus-based sensitivity in chromosome 1 and 3, but has slightly lower than find circ in chromosome 2 (Figure 5 B). Moreover, CircMarker only needs around 15 minutes to finish the whole analysis of teated sample while other tools may take at least 1 hour (CIRCexploprer even takes more than 9 hours). Overall, CircMarker outperforms the other tools on this data.

**Fig. 5.**
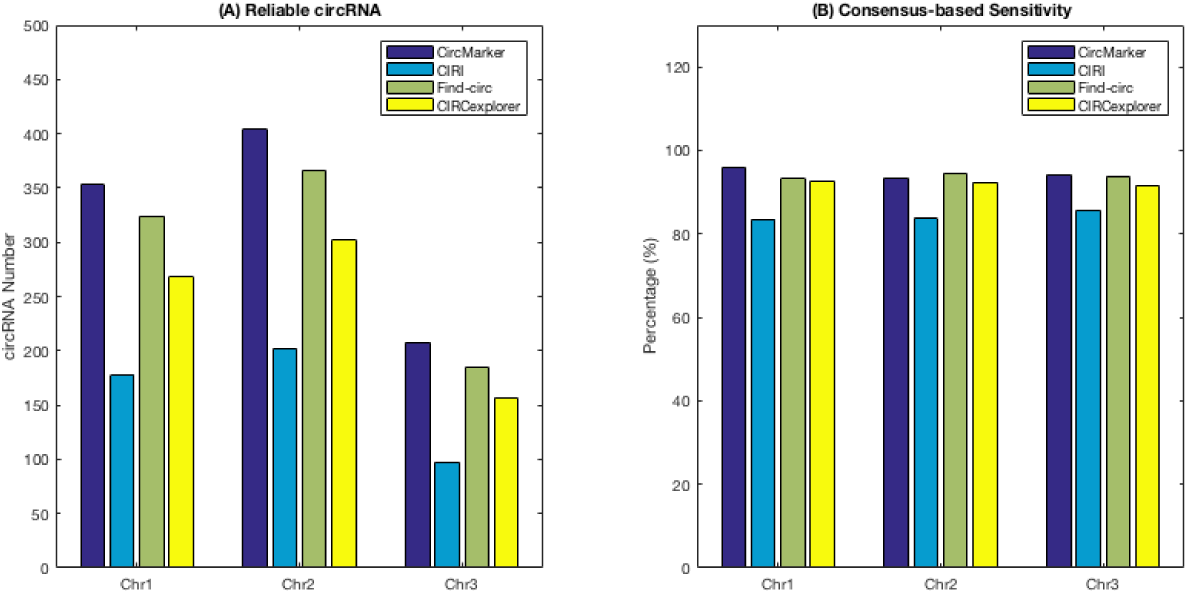
(A) The number of reliable circular RNAs called by each tool from chromosomes 1 to 3. The reliable circular RNAs come from the candidates which could be found in both treated and untreated sample. (B) The consensus-based sensitivity of each tool, which measures how many benchmark (i.e. found by at least two tools) circular RNA be contributed by the reliable circular RNA from each tool.

## 4 Conclusion

In this paper, we develop a new circular RNA detection method called CircMarker based on k-mer analysis. CircMarker runs much faster than other tools because it doesn’t perform reads mapping. Moreover, k-mers contain useful information about circular RNA detection. Our results on both simulation data and real data demonstrate that CircMarker can find more circular RNA. It has higher consensus-based sensitivity and high accuracy/reliable ratio compared with others. In addition, the circular RNAs called by CircMarker often contain most circular RNAs called by other tools in the real data we tested. This implies that CircMarker has low bias. CircMarker is easy for use. CircMarker is a stand-alone tool (implemented by C++) and does not depend on any third party tools. The source code and test data can be downloaded at: https://github.com/lxwgcool/CircMarker.

## Acknowledgement

This work was supported by grants IIS-0953563, IIS-1447711 and IIS-1526415 from US National Science Foundation to YW.

